# Epistasis detectably alters correlations between genomic sites in a narrow parameter window

**DOI:** 10.1101/570903

**Authors:** Gabriele Pedruzzi, Igor M. Rouzine

## Abstract

Different genomic sites evolve inter-dependently due to the combined action of epistasis, non-additive contributions of different loci to genome fitness, and physical linkage of different loci due to their common heritage. Both epistasis and linkage, partially compensated by recombination, cause correlations between allele frequencies at the loci (linkage disequilibrium, LD). The interaction and competition between epistasis and linkage are not fully understood, nor is their relative sensitivity to recombination. Modeling an adapting population in the presence of random mutation, natural selection, pairwise epistasis, and random genetic drift, we compare the contributions of epistasis and linkage. For this end, we use a panel of haplotype-based measures of LD and their various combinations calculated for epistatic and non-epistatic pairs separately. We compute the optimal percentages of detected and false positive pairs in a one-time sample of a population of moderate size. We demonstrate that true interacting pairs can be told apart in a sufficiently short genome within a narrow window of time and parameters. Outside of this parameter region, unless the population is extremely large, shared ancestry of individual sequences generates pervasive stochastic LD for non-interacting pairs masking true epistatic associations. In the presence of sufficiently strong recombination, linkage effects decrease faster than those of epistasis, and the detection of epistasis improves. We demonstrate that the epistasis component of locus association can be isolated, at a single time point, by averaging haplotype frequencies over multiple independent populations. These results demonstrate the existence of fundamental restrictions on the protocols for detecting true interactions in DNA sequence sets.

## Introduction

Epistasis is inter-dependence of fitness effects of mutations occurring at different loci caused by biological interactions between domains of proteins and between proteins and nucleic acids [1-4]. In biological systems, amino acids in proteins domains interact with each other. The resulting networks of interactions that include direct protein-protein binding and allosteric effects, shape the gene regulation and metabolic networks. Epistasis is a widespread property of biological networks [2, 5-8] and a subject of intense studies. The vital role it plays in the genetic evolution of populations and the heritability of complex traits is well established. The existing estimates indicate that the variation of an inherited trait across a population can only partially be explained by the additive contributions from the relevant alleles. On average, 70% of the inheritance may be due to epistasis or epigenetic effects [9]. Epistasis defines the evolutionary paths and creates fitness valleys, i.e., intermediate genetic variants with reduced fitness [10-12].

A crucial biological scenario is a viral population adapting to the abrupt changes in external conditions. Examples include the transmission to a new host, the invasion of a new organ, or the process of immune evasion or the development of drug resistance. Typically, virus adaptation consists of primary mutations followed by a cascade of several compensatory (helper) mutations [13-18]. These mutations help the adapting virus to pass through a fitness valley [11]. During this process, compensatory mutations rescue the replicative fitness of virus while preserving its resistant phenotype [13, 15, 19].

However, epistasis is not the only force causing inter-dependence in the evolution of genomic regions. The other dominant factor is the host of linkage effects existing between genomic regions that co-evolve in the same time frame and share the same ancestors [20, 21]. They include Fisher-Muller effect (clonal interference), genetic hitchhiking and genetic background effects, and Hill-Robertson interference between genetic drift and selection [21-23].

The other effect of linkage is a genetic association between loci, or linkage disequilibrium (LD). The effects of linkage on the evolution of a long genome in the presence of selection is well understood theoretically [12, 24-31]. The theory shows that linkage significantly slows adaptation many times, enhances accumulation of deleterious mutations, and changes the shape of the phylogenetic tree [32, 33]. The magnitude of linkage effects grows rapidly with the number of loci, *L*. Recombination partly offsets linkage effects and accelerates evolution [34-40] and competes with epistasis [41]. Epistasis has been shown to be potentially important for the evolution of recombination in a two-locus model [42, 43].

One consequence of linkage at large *L* is the strong interaction between the evolutionary trajectories of different sites that, depending on the case, can be both positive and negative. LD stemming from this interaction is easy to confuse with epistasis effects. Linkage effects become small only in populations that are exponentially large in the number of sites *L* [25]. Further, working with sequence data from real populations, it is often unclear how to discriminate the effects of shared ancestry from those of epistasis, and which of the two evolutionary forces dominates in each case (for a comprehensive review, see [1, 44, 45]). Therefore, despite of a considerable theoretical and experimental effort, detecting epistasis from genomic data remains a challenge.

In the present work, we offer an evolutionary explanation for the observed difficulty of the detection of epistasis from one-time data set. The idea is to generate mock data using a Monte-Carlo model of evolution and then try to discriminate between effects of linkage and epistasis. We use a panel of six pairwise LD measures to compare their distributions between epistatic and random pairs in a broad range of model parameters. We also use 3D and 2D maps of all possible combinations of LD measures and employ an optimization algorithm based on *a priori* knowledge to estimate the best, theoretically possible identification of epistatic pairs. As a result, we delineate the region of time and model parameters where the epistatic pairs can be detected against the linkage background. Finally, we investigate the role of recombination and the effects of averaging over multiple independently-evolving populations.

## Results

### Computer simulation of evolution

We consider a haploid population of *N* genomic sequences comprised of *L* sites, where *L* >> 1, and either a favorable or deleterious allele is present at each site. Evolution of the population between discrete generations is simulated using a Wright-Fisher model including the evolutionary factors of random mutation with the rate *µ* per site, random genetic drift, and natural selection, as described in *Methods*. Natural selection includes positive (antagonistic) epistatic interaction between selected pairs of deleterious alleles. A simple case of genomes with uniform selection coefficient *s*_0_ and uniform epistatic strength, *E*, is considered. We also assume that epistatic pairs are isolated, i.e., that each genomic site interacts with only one site. The initial population is randomized as it is done in virus passage experiments, with an average allelic frequency *f*_0_. In most of our work, we initially neglect the factor of recombination and primarily focus on asexual evolution, but lift this restriction in the end and explore broad parameter ranges. We aim to simulate the detection of epistatic pairs and identify the best conditions for detection theoretically.

### Measures of linkage disequilibrium (LD)

Various haplotype-based measures based on known haplotype frequencies have been proposed to characterize the allelic association between loci. We will list four measures, as follows.

#### Lewontin’s measure

A classical measure of statistical correlation between alleles at different loci has a form [46]

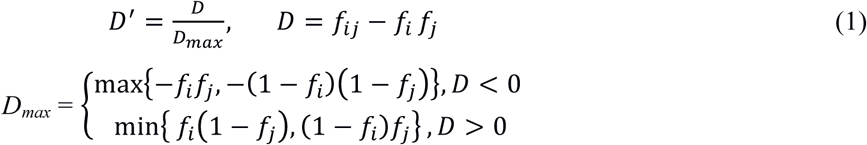

Here *f*_*ij*_ is the average frequency of a bi-allelic haplotype of loci *i* and *j*, and *D*_*max*_ is a normalization coefficient making sure that *D′* ∈ [0,1].

#### Pearson’s correlation coefficient

An alternative is the correlation coefficient between pairs of loci *r*, expressed as [47]

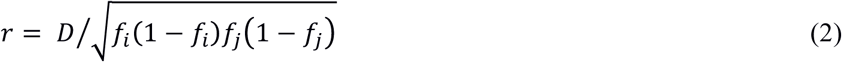

#### Kimura-Wu measure

More recently, Wu and colleagues proposed another statistical marker of linkage disequilibrium, which, for binary alleles, has a form [48]

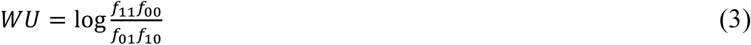

which represents the logarithm of the *Z*-measure proposed much earlier by Kimura [49].

#### Universal footprint of epistasi

In our recent work [50], we introduced another bi-allelic measure of LD

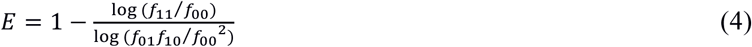

The advantage of this measure with respect to previous three is that it has a direct meaning in terms of fitness. For isolated interacting pairs, it represents the degree of mutual compensation of two deleterious mutations when frequencies in Eq. 4 are ensemble-averaged (see *Methods* below). Here the value *E* = 0 corresponds to the absence of compensation (epistasis), and *E* = 1 to full mutual compensation of the two mutations. Note the singularity in Eq. 4 at *f*_10_ *f*_01_ = *f*_00_^2^; we checked that it does not affect our results.

Below we investigate the effect of linkage for interacting and noninteracting pairs of loci using the measures defined in Eqs. 1-4. Also, we employ an optimization algorithm that, exploiting a priori knowledge of the correct epistatic pairs, puts the best possible threshold between the two distributions of LD. We consider different combinations of two or three LD measures to obtain the best detection possible.

### LD of epistatic and non-epistatic pairs are distinct in a narrow parameter window

We started by plotting the distribution of six LD measures calculated from Eq. 1 over individual pairs of sites, at different times (Fig. 1). We show separately the distribution for two subsets of pairs: the known epistatic subset (dark shade) and all the pairs (light shade). In the beginning, LD is narrowly distributed around zero, for both epistatic and non-epistatic subsets (Fig. 1, row 1).

**Fig. 1.**
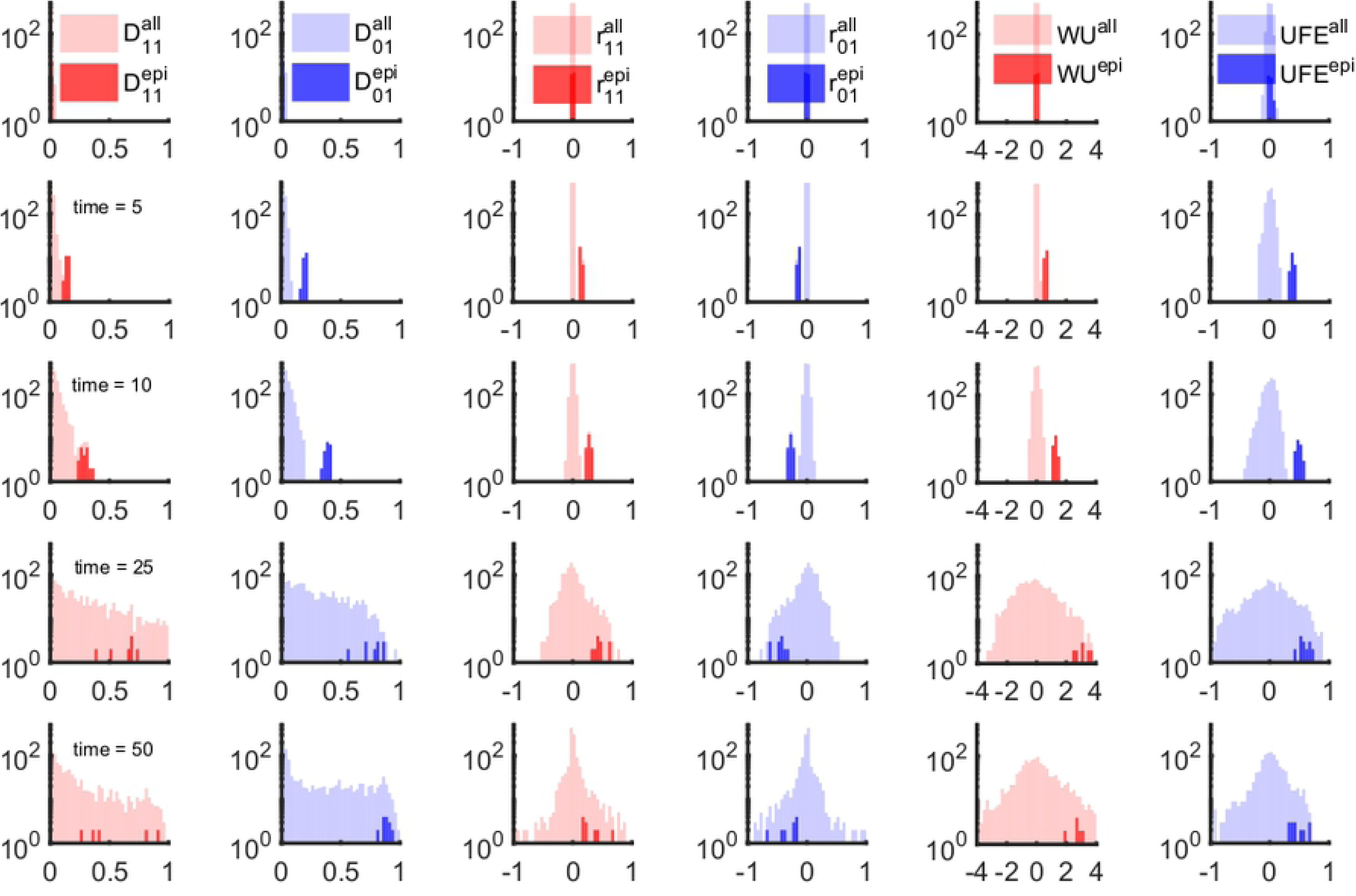
LD- and haplotype-based measures of epistasis identify a narrow time window of epistasis detectability. We compared the time-dependent distribution of 6 markers of LD shown in 6 columns. Each column show the profile of the distribution of a measure of epistasis: *D*_11_, *D*_01_ (Eq. 1), *r*_11_, *r*_01_ (Eq. 2), WU (Eq. 3) and UFE (Eq. 4). Different rows correspond to different time points: *t* = 1, *t* = 5, *t* = 10, *t* = 25 and *t* = 50. The shaded regions correspond to the density distributions for all possible pairwise interactions (lighter color) and the known epistatic pairs (darker shade). The shaded areas are normalized distributions reflecting the fact that epistatic pairs represent a tiny fraction of the all possible pairs in a genome. The fluctuations of non-epistatic pairs increasing in time overlap onto the distributions of epistatic pairs. Parameters: *N* = 2 10^4^, *s*_0_ = 0.1, *L* = 50, *E* in the range [0, 1], *µL* = 7 10^−2^. Each odd site interacts with its neighbor on the right (1-2, 3-4, 5-6,…) with epistatic strength *E* = 0.75. Initially, sequences were random with average allelic frequency set to *f* = 0.4. The negative control result in the absence of epistasis (*E* = 0) is presented on Supplementary Fig. S1.

Subsequent time points (Fig 1, rows 2 and 3) show progressive separation of the two distributions. In the course of further evolution (Fig. 1, rows 4 and 5), the distribution of randomly-chosen pairs, which was initially narrow and concentrated near the origin *E* = 0, gradually expands and overlaps with the small epistatic distrbution (Fig 1). This effect implies that non-epistatic pairs of sites, due to the stochastic nature of evolution, produce large LD of random sign. In this case, it is impossible to tell apart epistatic pairs from any of these measures of LD.

### Results are robust to the choice of an LD measure or their combination

Next, we checked whether combinations of LDs used together can improve detection. We have calculated all possible combination of six LD measures in Eq. 2 and tried to separate interacting and non-interacting pairs using 3D and 2D scatter plots. A representative example is shown in Fig. 2, for *E*=0, and for *E*=0.75 at two time points. Other possible combinations of 2 and 3 measures are summarized in Table S1 in *Supplement*.

**Fig. 2.**
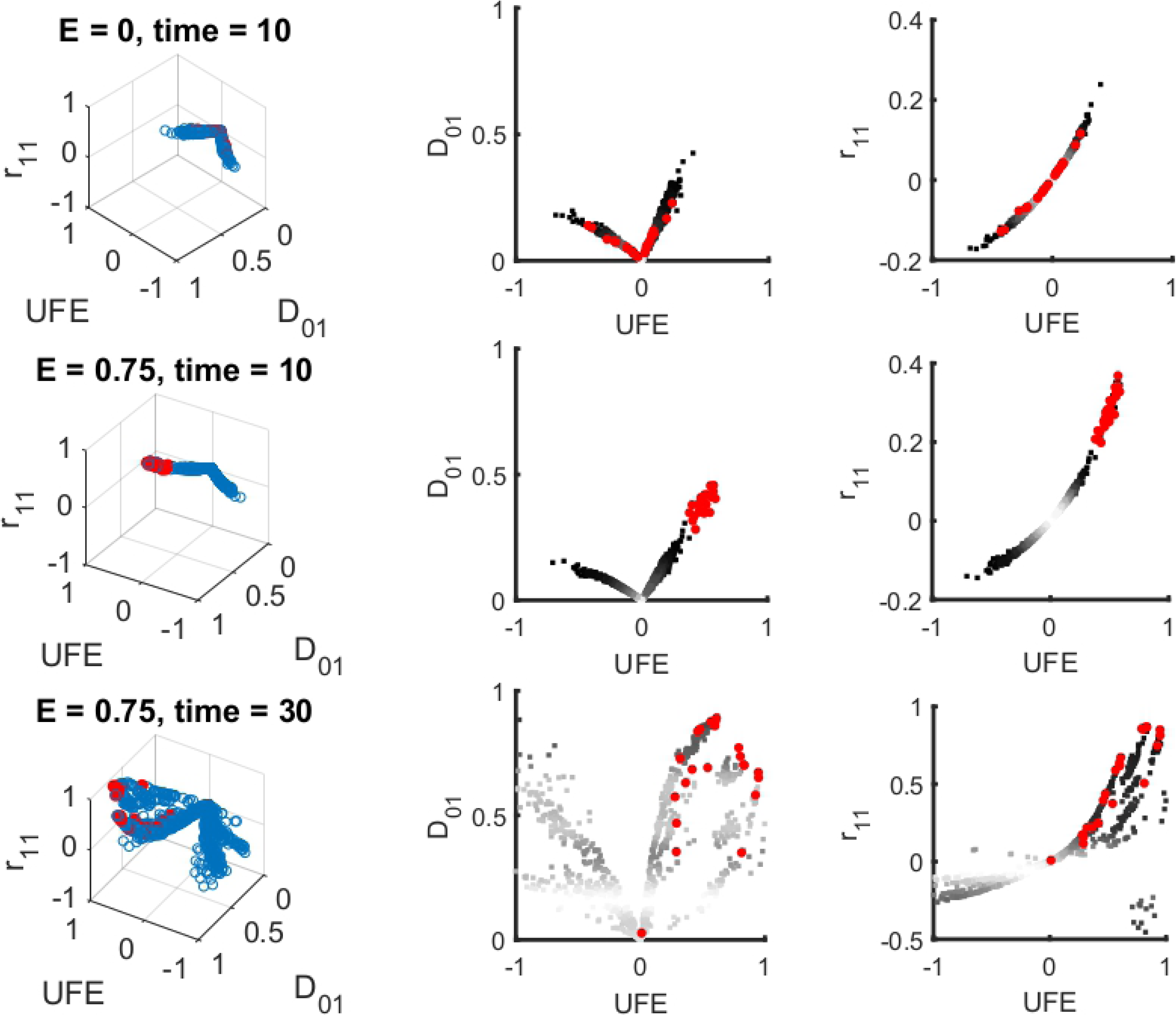
The optimization algorithm to identify ideal conditions for detection of epistasis is exemplified through the 3D scatter plot of three different measures of LD. Left: A representative example of three-dimensional scatter plots of three statistics, UFE, *D*’_01_ and *r*_*1*1_, plotted for all individual pairs of sites (blue circles) and for known epistatic pairs (red circles). Right and middle: two-dimensional projections. The three rows correspond to the absence of epistasis (E = 0, top), and two time points in the presence of epistasis, within the detection window and outside (middle and bottom). All possible combinations of two and three measures have been tested and summarized in Table S1. At intermediate time *t*=10, a district cloud of epistatic pairs (red dots) cluster together outside the overall distribution of all pairs and, hence are detectable. At long times, substantial overlap with non-interacting pairs bias contaminates detection. To optimize detection, we define a detection threshold for each of the detection variables (UFE, WU and the four haplotypes) and adopted an optimization algorithm that minimizes the following quantity “DET + *a* FPOS", where *a* is a fitting parameter, DET represent the detection percentage, and FPSO is the percentage of false positive, based on prior knowledge of the identity of true epistatic pairs. Parameters as in Fig. 1

We wrote an optimization algorithm which separates the cloud of interacting pairs from the cloud of non-interacting pairs in the best possible way, using *a priori* knowledge about the identity of pairs (Fig. 2). We adjusted the threshold to optimize the difference between the detection rare and the false positive rate. This method, employing the principle of machine learning, does not give any substantial improvement on the detection window (See Supplementary Table 1). For a real data sets, *a priori* knowledge about interacting pairs is usually unavailable, so that the detection of epistasis in a single population at one time point will be even worse than our prediction.

### Parameter sensitivity analysis confirms the narrow window of detection

#### Selection coefficient

Next, we investigated how the window of detection changes with model parameters. We calculated the detection rate and the false positive rate for the six measures of LD at different values of selection coefficient, *s*_0_ (Fig. 3). For each measure, the results show an inverse scaling of the detection time window on *s*_0_. Note that the window closes at very small *s*_0_, where evolution is almost selectively neutral, and epistasis is never detectable.

**Fig. 3.**
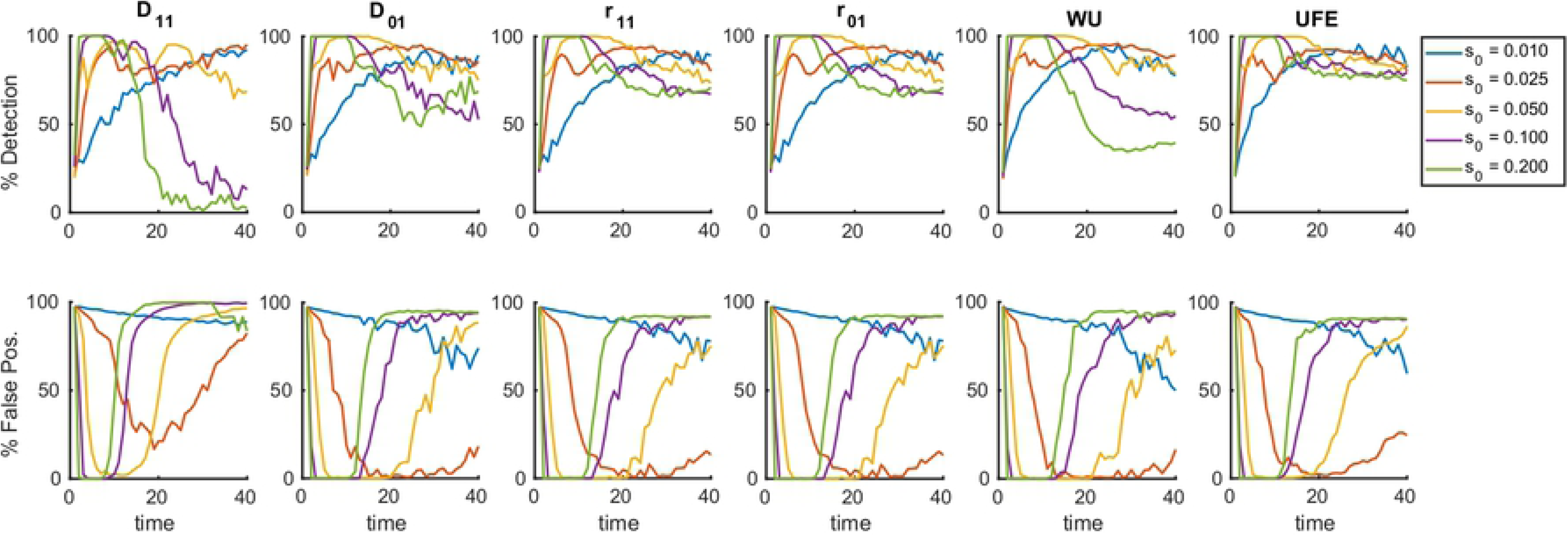
Detection of epistasis is confined in a time window whose width is controlled by the mean selection coefficient. Percentile of detection and false discovery as a function of time is averaged over 25 random simulation runs per each value of *s*_0_, the constant selection coefficient for each allele in the sub-population. The detection of epistatic pairs for a panel of measures of LD, namely, *D*_11_, *D*_01_ (Eq. 1), *r*_11_, *r*_01_ (Eq. 2), WU (Eq. 3) and UFE (Eq. 4). Results from a detection protocol that maximizes the difference between the detection percentile and the false-positive fractions by tuning the detection threshold, show the same trend for all measures considered. At time ∼1.5/*s*_0_ generations, we observe the beginning of a transition which completely blurs the detection of epistatic interaction at time ∼2.5/*s*_0_. The initial allelic frequency *f*_0_ = 0.45, *s*_0_ is shown, the other parameters are as in Fig. 1.

#### Distributed selection coefficien

Next, we conducted a sensitivity analysis with respect to the other model parameters (Fig. S5). Firstly, we lifted the simplifying assumption of a constant selection coefficient, *s*= *s*_0_, and allowed variation of *s* among sites according to a half-Gaussian distribution. We obtain a similar dependence of the window width on the average selection coefficient (Fig. S5), although with a higher false positive rate within the detection window than for the case with constant *s*.

#### Length of the genome

We found out, that sequence length *L* limits the detectability of epistasis substantially (Fig. S5). An increase of the sequence length or a reduction of the population size leads to narrowing and, eventually, disappearance of the detection window. These results limit the applicability of these methods to short sequences. Indeed, the number of all possible locus pairs increases with genome length *L* proportionally to *L*^2^, and the number of epistatic pairs increases only as *L*, so that the task of finding “the ruby in the rubbish” becomes harder at larger *L* [1, 44, 45].

#### Population size

We observed a very slow (logarithmic) expansion of the detection window with population size *N* (Fig. S5). This is consistent with the results of asexual evolution models, which predict a very slow logarithmic dependence on *N* for all the evolutionary observables, including evolution speed, genetic diversity, and the average time to most recent ancestor [25-31, 35-37, 39, 40, 51]. Only in very large populations whose size increases exponentially genome length *L*, linkage effects become small [25]. In these, astronomically large populations, epistasis would be easily detectable.

#### Initial standing variation

We have observed a detection window in time only at the initial frequencies of deleterious alleles above 10% (Fig. S5). At smaller frequencies, detection lapses. We can conclude that detection of epistasis in a single population studied is possible in a narrow parameter range.

### Recombination improves detection

Until now, we have assumed a completely asexual evolution. In our next step, we investigated the role of recombination, parametrised by the average number of crossovers per genome, *M*, and by the probability of outcrossing per genome, *r*. We obtained that intermediate recombination rates rescue the detection of epistasis by disrupting linkage and yet preserving the epistasis contribution to LD. At our default parameter set (Fig. 1), we observed a significant reduction of linkage fluctuations starting from *r* = 20% and *M* = 5 (Fig. 4). The results show that LD effects of linkage are much more resistant to recombination than, for example, the evolution speed, which increases substantially already at tiny values of *r* [34-40]. We found out also that extremely high levels of recombination decrease LD for epistatic pairs as well, thus rendering epistasis undetectable. Thus, there exists a narrow window of recombination rates where epistasis can be observed outside of the detection window for time and other parameters described above.

**Fig. 4.**
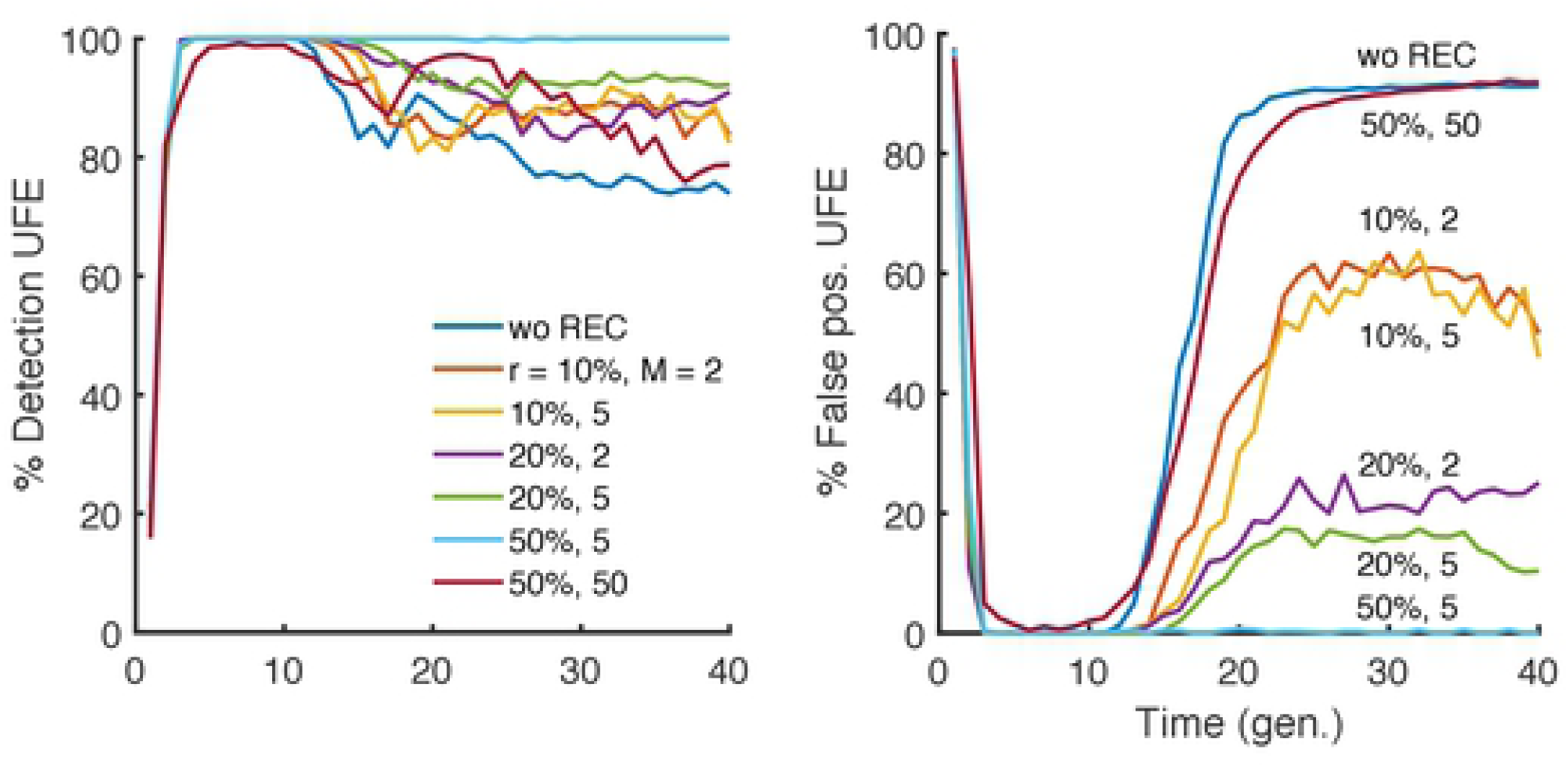
Variation of the time window of detection with recombination. Percentile of detection and false discovery as a function of time is averaged over 25 random simulations (runs) in a broad range of parameters values. The detection rare and false positive rate of epistatic pairs with UFE at different values of *s*, randomly drawn from a half-Gaussian distribution of deleterious alleles. The presence of moderate recombination characterized by outcrossing rate *r* and the average number of cross-overs, *M*, broadens the detection window. We observe similar results for all the statistics considered in this study (data not shown). The default parameter set is *E* = 0.75, with the other parameters as in Fig. 1.

### Population divergence creates strong linkage effects

In order to understand the reason behind the strong linkage effects masking epistasis, we investigated the time-dependent changes of the phylogenetic tree using a hierarchical clustering algorithm (Fig. 5a-d). The initial, randomized population display a star-shaped phylogeny, characterized by the same mean distance between all sequences and the most common sequence (Fig. 5). With time, the phylogenetic tree grows branches of increasingly related sequences (Fig. 5c, d). As simulation continues (Fig. 5d), the tree becomes more lopsided, while recent mutations create short branches at the bottom. At the same time, we observe that the tree has a decreasing number of ancestors. Eventually, the tree evolves into Bolthausen-Sznitman coalescent (BSC) with a single common ancestor, previously predicted for the stationary regime of traveling wave [25, 29, 37] (Fig. 5).

**Fig. 5.**
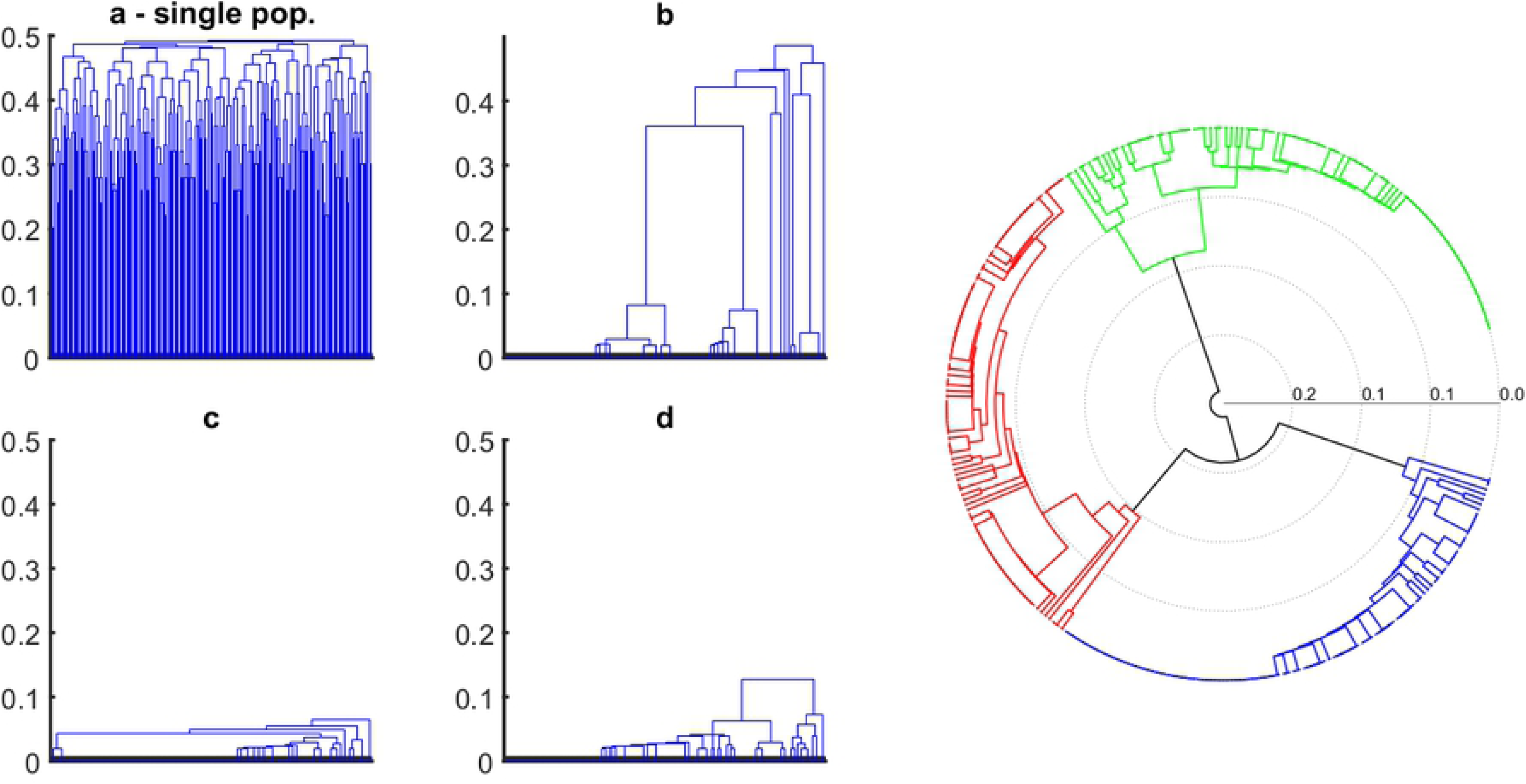
Evolution of genealogy within a single, well-mixed population and comparative representation of multiple, independently evolving population. (a-d) Phylogenetic structure of a single population comprising a sample of 500 genomes at four different times: *t* = 0, 10, 20, and 30 generations. Mean genetic distance between genomes decreases in time, and the structure of the tree changes from a star-like shape towards a monophyletic tree (BS coalescent), with a single common ancestor. The right panel shows the reconstructed phylogenetic tree of three populations, independently evolved from the same initial random seed. At a glance, it is possible to determine that the three populations do not share much sequence homology and segregate into different, phylogenetically distinct clades. *N*= 20000 genomes, initial average allelic frequency *f*_0_ = 0.40, other parameters as in Fig. 1.

Emergence of this phylogeny is coincident with the increase in the fluctuations of LD of non interacting pairs (Fig. 1). The reason for strong random LD is stochastic divergence of the population from the initial state, as illustrated by clustering of three independently evolved populations (Fig. 5, right). The distance between the trees obtained in separate runs increases linearly in time due to fixed beneficial mutations at randomly chosen sites. Haplotype configurations of the common ancestor of the population are inherited by all members of the population, with some small variation determined by the time to the most recent common ancestor. Thus, the stochastic divergence of individual populations creates strong LD with a random sign.

### The use of multiple populations defeats LD fluctuations and rescues epistatic signature

Because the linkage fluctuations arise due to stochastic divergence of the founder, the common ancestor, the natural idea is to use multiple populations to average over possible founder sequences. To test this idea, we evolved independently multiple populations at the same initial conditions and averaged the haplotype frequencies used in LD markers (Eqs. 1-4) over populations, for each pair of sites, separately. We found out that including a sufficient number of independent populations results in a substantial reduction of the noise and indefinite expansion of the window of detection (Fig. 6). Qualitatively similar results are obtained for all LD markers.

**Fig. 6.**
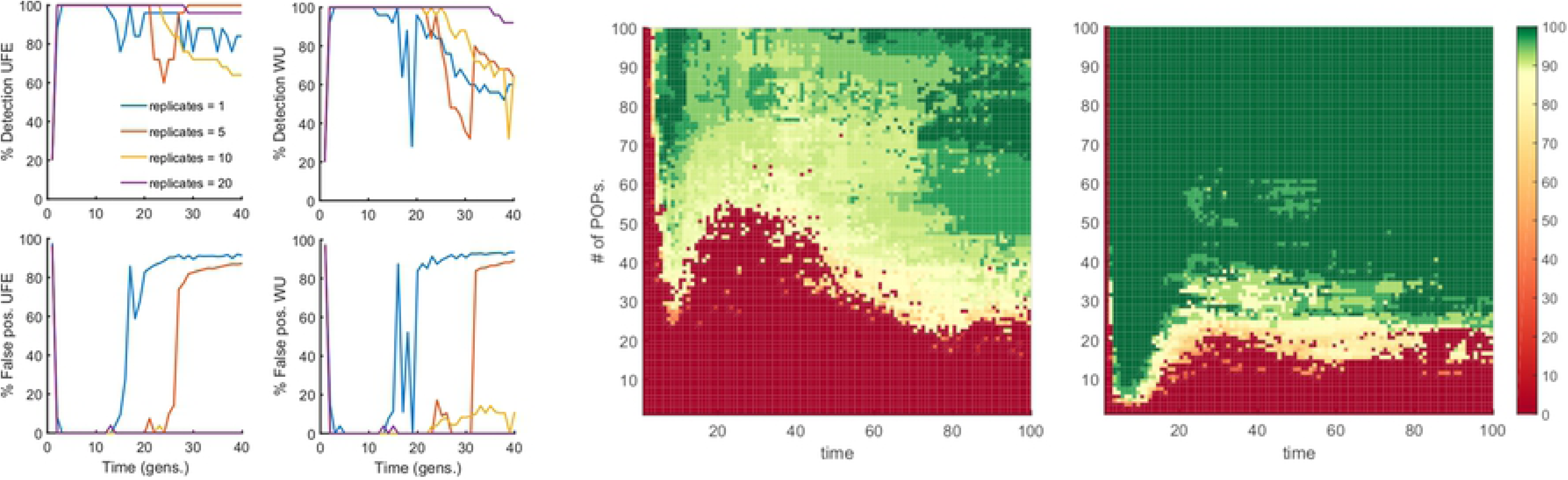
Detection of epistasis is rescued by simultaneous analysis of multiple independently-evolved populations. Left 4 plots: Percentile of detection (top)_ and false discovery (bottom) as a function of time are presented for UFE and WU measures. Number of replicate Monte-Carlo runs is shown. The haplotype frequencies are averaged over runs, which represent independently-evolved populations. At time ∼1.5/*s*_0_, we observe the beginning of a transition which completely blurs the detection of epistatic interaction for a single replicate (blue line), however, already 5 replicates are sufficient to significantly extend the detection window up to ∼2.5/*s*_0_, and a higher number of replicates completely eliminate false-positive pairs, while maintaining the average detection above 80%. Parameters: *E* = 0.75, *N* = 20000, the others as in Fig 1. Right: Two-dimensional color maps for UFE measure of LD, which summarize the results of a similar analysis for two population sizes: *N* = 100 (middle plot) and *N* = 1000 (right plot). *Y*-axis: Number of independent populations. *X*-axis: time of evolution. Color shows the percentage of detection with the detection threshold of interacting pairs chosen to give the false discovery rate below 20%.

## DISCUSSION

In the present work, using a Monte-Carlo simulation of a haploid population, we calculated the distributions of six measures of linkage disequilibrium and their combinations for epistatic and random locus pairs. We demonstrated that, in a single asexual population, the footprints of epistatic pairs are readable only in a narrow time interval between 0.2/*s*_0_ and 1.5/*s*_0_ generations. During later adaptation, the distribution of linkage disequilibrium for non-interacting pairs broadens and engulfs the distribution for epistatic pairs. These results indicate that, long before the onset of the steady state, linkage effects dominate over the effects of epistasis. This phenomenon is predicted in a broad parameter region and for all the LD statistics, suggesting that, in the context of inherited linkage fluctuations, all statistics based on pairwise linkage disequilibrium are equal.

To gain insight into the evolutionary origin of these fluctuations, we investigated phylogenetic trees of the entire population at different time points to observe that the shape of the tree strongly correlates with the magnitude of linkage fluctuations. The shape of the phylogenetic tree changes in time from the initially star-shaped genealogy to a Bolthausen-Sznitman (BS) coalescent [32, 33] previously analyzed in great detail for adapting asexual populations [25, 36, 37]. Once BS genealogy is established, individual sequences share a high degree of interrelatedness due to fixed beneficial mutations at randomly chosen sites. The presence of the BS coalescent is coincident with strong co-inheritance linkage fluctuations. The stochastic nature of their common ancestor sequence, divergent in time from common ancestors in other independent populations (Fig. 5) is the cause of the strong fluctuations of LD.

We have also directly quantitated the detection of epistatic pairs against the background of random linkage effects. We evaluated the sensitivity of the width of the detection window with respect to several input parameters, such as the mean selection coefficient, the size of the population, the sequence length, and initial genetic variation, and the role of recombination. We observed that the window is proportional to the inverse average selection coefficient, 1/*s*_0_, but a very small *s*_0_ abolishes any chance of detection, so that the best detection is attained in the case of moderately weak selection. The detection window exists only for sufficiently small genomes. The presence of recombination has the effect of compensating the linkage component and thus significantly improving the detection of epistasis. Yet, very frequent recombination disrupts epistatic effects.

To isolate the epistatic component from co-inheritance effects, we performed simulations over several independently-evolved populations and averaged the haplotype frequencies over these runs. The results predict the number of independent population required to attain significant expansion of the detection window (Fig. 6). Thus, the averaging over multiple independently-evolved populations filters out linkage effects leaving a clear footprint of epistasis in a much broader parameter range. However one should note that the multiple-population sampling was conducted under the ideal conditions, in which every population evolved independently for the same time with the same parameter set, and represented the same fraction of the total sample. Unequal sampling or heterogeneous representation in real data sets may create additional problems.

Our model adopts several simplifying assumptions. (i) Deleterious alleles are assigned selection coefficient constant in time. (ii) We considered constant and fixed epistatic strength for all pairs. (iii) We focused on a simple topology of epistatic network. While these are reasonable assumptions to describe the problem of linkage fluctuations in biological systems, a real scenario with mixed sign epistasis and complex topology might pose additional challenges for the accurate detection of epistasis.

In summary, we offer an evolutionary reason for the fluctuations of epistatic estimates in the existing sequence sets. Linkage due to stochastic divergence of the common ancestor of a population from the origin is responsible for the high false-positive rates of epistasis detection in a single population. We demonstrated how the use of multiple independently-evolving populations, or the use of time series when available, allows us to average out strong linkage effects and rescue the detectability of epistasis.

## METHODS

We consider a haploid population of *N* binary sequences, where each genome site (nucleotide position) numbered by *i* =1, 2,…, *L* is either *K*_*i*_ =0 or *K*_*i*_ =1. We assume that the genome is long, *L* >> 1. Evolution of the population in discrete time measured in generations is simulated using a standard Wright-Fisher model, which includes the factors of random mutation with rate *μL* per genome, natural selection, and random genetic drift. Recombination is assumed to be absent. Once per generation, each genome is replaced by a random number of its progeny which obeys multinomial distribution. The total population stays constant with the use of the broken-stick algorithm. To include natural selection, we calculate fitness (average progeny number) *e*^*W*^ of sequence *K*_*i*_ as given by [50]

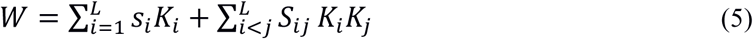

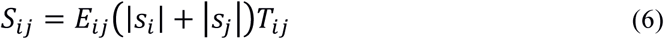

The first term in Eq. 5 stands for the additive contribution of single mutations to fitness with selection coefficients *s*_i_. The second term in Eq. 5 describes pairwise interactions of sites with magnitudes *S*_*ij*_, which are given by Eq. 6. Coefficient *E*_*ij*_ represents the relative strength of epistatic interaction between sites *i* and *j*, while the binary elements of the matrix **T** indicate the interacting pairs by *T*_*ij*_ = 1 and the other pairs by *T*_*ij*_ = 0. An example of positive epistasis is the compensation of two deleterious mutations inside protein segments that bind each other. Note that *E*_*ij*_ = 1 corresponds to full mutual compensation of deleterious mutants at sites *i* and *j*. We consider the simplest interaction topology of interacting neighbors, as given by *T*_2*i*,2*i*+1_ = 1 and 0 for all other pairs.

## Acknowledgments

We thank Martin Weight for insightful comments.

